# Single-strand DNA processing: phylogenomics and sequence diversity of a superfamily of potential prokaryotic HuH endonucleases

**DOI:** 10.1101/279083

**Authors:** Yves Quentin, Patricia Siguier, Mick Chandler, Gwennaele Fichant

## Abstract

**Background:** Some mobile genetic elements target the lagging strand template during DNA replication. Bacterial examples are insertion sequences IS*608* and IS*Dra2* (IS*200*/IS*605* family members). They use obligatory single-stranded circular DNA intermediates for excision and insertion and encode a transposase, TnpA_IS*200*_, which recognizes subterminal secondary structures at the insertion sequence ends. Similar secondary structures, Repeated Extragenic Palindromes (REP), are present in many bacterial genomes. TnpA_IS*200*_-related proteins, TnpA_REP_, have been identified and could be responsible for REP sequence proliferation. These proteins share a conserved HuH/Tyrosine core domain responsible for catalysis and are involved in processes of ssDNA cleavage and ligation. Our goal is to characterize the diversity of these proteins collectively referred as the TnpA_Y1_ family.

**Results:** A genome-wide analysis of sequences similar to TnpA_IS*200*_ and TnpA_REP_ in prokaryotes revealed a large number of family members with a wide taxonomic distribution. These can be arranged into three distinct classes and 12 subclasses based on sequence similarity. One subclass includes sequences similar to TnpA_IS*200*_. Proteins from other subclasses are not associated with typical insertion sequence features. These are characterized by specific additional domains possibly involved in protein/DNA or protein/protein interactions. Their genes are found in more than 25% of species analyzed. They exhibit a patchy taxonomic distribution consistent with dissemination by horizontal gene transfers followed by loss. The *tnpA*_REP_ genes of five subclasses are flanked by typical REP sequences in a REPtron-like arrangement. Four distinct REP types were characterized with a subclass specific distribution. Other subclasses are not associated with REP sequences but have a large conserved domain located in C-terminal end of their sequence. This unexpected diversity suggests that, while most likely involved in processing single-strand DNA, proteins from different subfamilies may play a number of different roles.

**Conclusions:** We established a detailed classification of TnpA_Y1_ proteins, consolidated by the analysis of the conserved core domains and the characterization of additional domains. The data obtained illustrate the unexpected diversity of the TnpA_Y1_ family and provide a strong framework for future evolutionary and functional studies. By their potential function in ssDNA edition, they may confer adaptive responses to host cell physiology and metabolism.

## Background

HuH enzymes are dedicated to processing single-strand DNA (ssDNA) and use particular DNA recognition and reaction mechanisms for site-specific ssDNA cleavage and ligation. Members of this protein family are numerous and widespread in all three domains of life. There are two major classes within the HuH superfamily [1]: the Rep (replication) proteins and the relaxase or Mob (mobilization) proteins which process DNA during plasmid replication and conjugation, respectively. However, HuH endonucleases have also been identified in other processes involving ssDNA, such as replication of certain phages [2] and eukaryotic viruses [3], and in different types of transposon. These proteins have also been appropriated for cellular processes such as intron homing [4] and processing of bacterial <u>R</u>epetitive <u>E</u>xtragenic <u>P</u>alindromic (REP) sequences [5]. The family relationship is based on several conserved amino acid motifs. These include the HuH motif [6,7], composed of two His residues (H) separated by a bulky hydrophobic residue (U), and the Y motif, containing either one or two Tyr (Y) residues separated by several amino acids. Together, these constitute the core catalytic domain.

Many family members include additional functional domains such as helicases, primases or zinc fingers. The simplest examples are transposases of the IS*200*/IS*605* insertion sequence (IS) family (TnpA_IS*200*_) [8]. They are approximately 150 amino acids long, possess only the HuH-Y core domain of about 113 amino acids (Figure 1) and function as dimers [9,10] (Figure 1A). The two H residues provide two of the three ligands involved in coordinating an essential Mg^2+^ ion. The third amino acid involved in this is located approximately four residues downstream from the catalytic Y residue (Q131 in the *Helicobacter pylori* IS608 transposase and N119 in TnpA_REP_ from *Escherichia coli* MG1655, Figure 1D). The Y residues act as nucleophiles in the cleavage reactions generating covalent phosphotyrosine intermediates. They use ssDNA IS substrates and catalyze IS insertion into and excision from the lagging strand template at replication forks [11] via a circular ssDNA intermediate. The HuH transposases, TnpA_IS*608*_ and TnpA_IS*Dra2*_ (from IS*608* and IS*Dra2* respectively) have been extensively studied at the functional level using a combination of genetics, biochemistry and structural biology [9,10,12–21] and the transposition pathway has been described in detail [11]. These enzymes do not directly recognize the DNA sequences which they cleave at each IS end during transposition. Instead, they bind small subterminal DNA hairpin structures of about 20-25 nt at both the left (LE) and right (RE) ends. The cleavage position is determined by a complex series of base interactions between a tetranucleotide (guide sequence) just 5’ to the hairpin foot and a conserved target tetranucleotide which flanks LE. At the right end, similar types of interaction occur between the final tetranucleotide of RE and an equivalent tetranucleotide guide sequence. The interactions can involve bases which form canonical (Watson and Crick) and non-canonical interactions including base triples [10,14].

**Figure 1:**
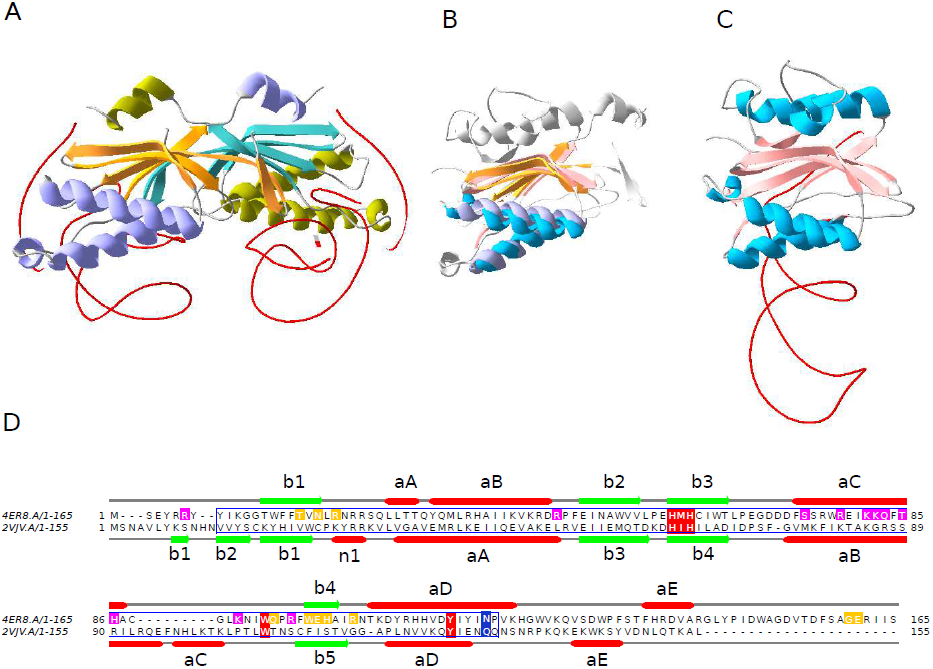
Structural homology between TnpA_IS*200*_ and TnpA_REP_. Panel A, TnpA_IS*200*_ from *H. pylori* TnpA_IS*608*_ dimer (PDB 2VJV) as a ribbon diagram with one molecule colored cyan (β sheets) and purple (α helix) the other molecule in orange (β sheets) and green (α helix), with the DNA in red. Panel C, *E. coli* TnpA_REP_ monomer (PDB 4ER8). Panel B, TnpA_IS*608*_ and TnpA_REP_ monomer structure superposition, with un-conserved structural elements in grey. The superpositioning was accomplished using SWISS-MODEL and the visualization with Swiss-PdbViewer [86](http://www.expasy.org/spdbv/). Panel D, TnpA_REP_ (4ER8) and TnpA_IS*608*_ (2VJV) sequence alignment with the secondary structures. HuH, W and Y conserved residues in red boxes, the residues essential for transposition in vivo (Q131 in *H. pylori* and N119 in TnpA_REP_ from *E. coli* MG1655) in blue, residues involved in 5’ GTAG interaction in yellow and residues REP sequence hairpin interaction in magenta, the core conserved domain in blue frame.

Closely related members of this group are the REP-associated tyrosine transposases, RAYTs [22] also referred as TnpA_REP_ to highlight their functional proximity with TnpA_IS*200*_ [5,22]. These also include the HuH-Y functional core [23]. In contrast to TnpA_IS*200*_ (Figure 1A), TnpA_REP_ from *E. coli* appears as a monomer (Figure 1C). The proteins share significant structural similarities (Figure 1B).

REP sequences provide genomic binding sites for proteins such as Integration Host Factor, DNA polymerase I, and DNA gyrase [24–26], can increase mRNA stability [27] and cause transcription termination [28,29]. They have also been implicated in regulating translation [30]. In many ways REP sequences resemble the subterminal secondary structures at the ends of IS*200*/IS*605* family members and are of similar size (21-65 nt). Like the IS LE and RE, they also carry a tetranucleotide guide sequence 5’ to the foot of the REP sequence hairpin necessary for cleavage by TnpA_REP_ [5]. As their name suggests, they are found largely between genes and are often present in many copies in a given host genome (up to 1% of *E. coli* chromosomes). They can be present as isolated copies or in clusters and often form particular structures, BIMES (bacterial interspersed mosaic elements [31]), whose basic unit includes two REP sequence copies in inverted orientation. These were later renamed REPINs (REP doublets forming hairpINs; [32,33]). There are a number of different REP sequence families or groups and several different groups can sometimes be found within a single genome [22,34–37]. In general, there appears to be only a single full length *tnpA*_REP_ gene for each REP sequence type and this is often flanked by a number of REP sequences forming a structure called a REPtron [5]. REPtrons do not appear to transpose as a unit but the REPs/BIMEs are likely to be mobilized by TnpA_REP_ activity [5,22,32,33]. Indeed TnpA_REP_ is able to cleave and rejoin REP sequences [5].

The apparent stability of *tnpA*_REP_ genes raises the question of how and why they are maintained in their host bacterial genomes. This was recently addressed through an analysis of *tnpA*_REP_ properties and a comparison to transposases of IS families (IS*200*/IS*605* and the entirely unrelated IS*110*) and two housekeeping genes [38]. The observations that *tnpA*_REP_ share more characteristics with housekeeping genes than with insertion sequences and that *tnpA*_REP_ are predominantly vertically transmitted, at the subgenus level, suggest that they have a yet uncharacterized function(s) that benefit their host cell and hence insure their maintenance [38].

Our goal in the work presented here was to characterize the diversity of this large protein family referred to collectively as the TnpA_Y1_ family, since all members share the conserved HuH/Y core domain responsible for ssDNA cleavage and ligation [11]. We provide a detailed classification of family members which was consolidated by the analysis of the conserved core domains and the characterization of additional domains. Only one subclass has the characteristic of insertion sequence transposases while five subclasses are more closely related to TnpA_REP_ proteins and are flanked by typical REP sequences in a REPtron-like arrangement. Other subclasses are not associated with REP sequences and illustrate the unexpected diversity of proteins belonging to the TnpA_Y1_ family. The data obtained provide a strong framework for future evolutionary study and the characteristics describing each subclass may help to further experimentally unravel their functions which are still largely unknown.

## Results & discussion

### Overall class and subclass organization of the TnpA_Y1_ family

Our initial TnpA_Y1_ library was composed of 924 proteins (Materials and Methods). To explore the family organization, the graph of the large connected component used in constructing the library (Materials and Methods) was reprocessed with MCL. The inflate factor (IF) value is an important parameter of MCL as it regulates the cluster granularity. We tested different IF values and analyzed their impact on graph partitioning. An IF of 1.2 generated three distinct classes (Figure 2B). Class 1 includes the TnpA_IS*200*_; class 2, the TnpA_REP;_ while the smaller class 3 contains a previously unknown family of HuH-Y proteins. Increasing IF values increased the number of sub-groups (Figure 2A) from five (IF values of 1.4) to thirteen (IF values > 3.8). This increase resulted from hierarchical decomposition of class 1 and class 2 into subclasses. Class 3 remained coherent and did not decompose into subclasses. We chose an intermediate IF value of 3.0 since the number of partitions remained stable at 12 subclasses over the range 3.0 ≤ IF ≤ 3.6. Here, class 1 forms two subclasses, 1.1 and 1.2, class 2 forms nine subclasses, 2.1 to 2.9, (Figure 2C), while class 3 remains unified. This classification into 12 subclasses was subsequently propagated to all proteins of the sample to obtain a complete overview (*i.e.* sequences of each cluster received the subclass of its medoid; Materials and Methods). 4,043 of the 4,081 initial proteins fell into the 12 TnpA_Y1_ subclasses. The remaining 38 proteins were false positives and were discarded.

**Figure 2:**
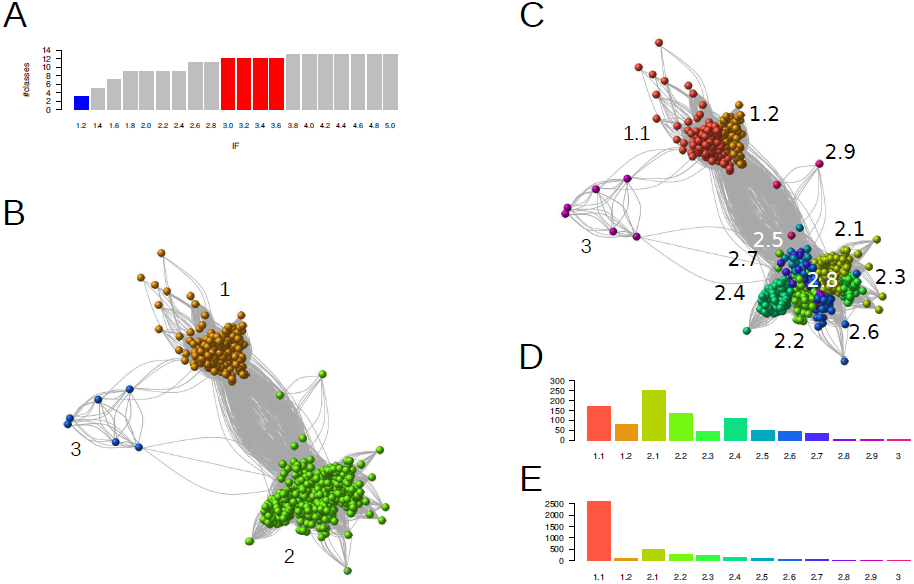
Family and sub-family organization of the 931 TnpA_Y1_ protein candidates explored with MCL. A. Evolution of the number of classes as a function of IF value. Three classes (family level) are obtained with IF=1.2 (blue bar) and rise to reach a maximum of 14 classes. 12 classes (subclass level) were obtained with IF in the range [3.0, 3.6] (red bar). B. Graph partitioning obtained with IF=1.2. Edges are BlastP hits (e-value < 1e-5) between proteins (vertices). The vertices are colored according to MCL classes. C. Graph partitioning obtained with IF=3.0. D. Histograms of the subclass sizes obtained with IF=3.0. E. Histograms of the subclass sizes obtained after propagation of the annotation to the complete sample of protein candidates. A maximum of 2,620 proteins is observed in subclass 1.1. The color code is the same in C, D and E panels.

The distribution of sequences into subclasses was uneven both for the initial (Figure 2D) and the full protein sets (Figure 2E). In both, there is a preponderance of subclass 1.1 (TnpA_IS*200*_) (171 in the restricted sample and 2,620 in the complete set due to multiple genomic copies of IS*200*/IS*605* family members). The contribution of the second most abundant class, 2.1, decreased in the full protein set.

An additional filter was included based on protein length within each subclass. The median protein length varies from 148 to 325 AA between different subclasses (Figure S1). Subclass 2.9 included only three proteins and was eliminated from further analysis. A few unexpectedly long proteins, the result of gene fusions or annotation errors, observed in each subclass and a number of significantly shorter sequences (partial) observed in almost all subclasses were also discarded (Figure S1). Partial sequences were significantly more abundant in subclass 1.1 (TnpA_IS*200*_) which show a tendency to decay [11].

### Subclass characterization

#### Conserved core domain

Class 2 members, which include TnpA_REP_ proteins, were initially retrieved with PF01797 (121 amino acids long, http://pfam.xfam.org/family/Y1_Tnp) which was built uniquely for the IS*200*/IS*605* family. However, the profile of most class 2 subfamilies aligns only partially with TnpA_REP_ sequences (figure S2A) and in some cases, two alignments are obtained for the N- and C-ter regions respectively. This is also true if the length of the aligned part of the protein with the profile is analyzed (Figure S2B).

For better core domain coverage, we constructed a specific HMMER TnpA_REP_-like family profile (Materials and Methods). The multiple alignments were manually edited to remove highly divergent sequences and to extract the conserved core domain as defined in PF01797. The TnpA_REP_-like profile obtained with hmmbuild from 428 aligned sequences is 115 amino acids long. This new profile allows excellent coverage of class 2 sequences: a complete alignment of the profile with the protein sequences is observed for all the subfamilies (Figure S2C) despite the sequence length variability of class 2 proteins (Figure S2D).

Class 3 includes only 10 members, 5 of which do not have conserved HuH or Y motifs. To augment this number, we searched Genbank for other instances of this class using BlastP. A sample of 24 sequences was retrieved and used to construct a specific class 3 profile with HMMER. Compared to class 1 and 2, class 3 proteins have an extended sequence conservation in the N-ter region leading to a profile length of 155 AA.

#### HuH and Y motifs

An *ab initio* search for conserved motifs with MEME confirmed the conservation of subclass-specific motifs overlapping the ubiquitous HuH, Y together with a W residue (Figure 1D and Figure S3). The distance between HuH and Y motifs is relatively constant (median between 58 and 67 AA) in the majority of subclasses, but is larger in subclasses 2.3, 2.4 and 2.6 (median 110, 88 and 82 amino acids) while subclass 2.7 and class 3 have a more disperse distribution (Figure S4). MEME also revealed additional short conserved motifs upstream (subclasses 2.4, 2.5, 2.7 and 2.8) and/or downstream (subclasses 2.1, downstream motifs share conserved serine residues and are part of a larger conserved domain (see below). The absence of conserved core motifs in certain sequences suggests that these correspond to non-functional proteins. To retain proteins that are most likely to be functionally active, we selected those that included a complete core domain defined by PF01797 or TnpA_REP_-like profiles encoding both the HuH and Y motifs. We obtained a sample of 722 from the original 924 sequences: 172 class 1, 545 class 2 and 5 class 3. In spite of the variations from class to class, the conservation of key catalytic HuH and Y residues suggest that these proteins are all involved in cleavage and rejoining ssDNA [1].

### Phylogenetic relationships between MCL classes

To gain insight into the relationship between these proteins, multiple alignments of the core domains of each class were first computed to take into account the variability in sequence length observed between MCL classes. These alignments were themselves aligned to produce a multiple alignment of the entire sequence set (Materials and Methods). This was trimmed to remove sites with rare indels and four unexpectedly divergent sequences. The final edited alignment contained 718 sequences with 121 sites. A phylogenetic tree (Materials and Methods) was computed and rooted on the branch that separates class 1 and 2 (Figure 3). We observed a strong correspondence between MCL subclasses and the tree topology although the former used complete sequences while the latter was based only on the core domain. Most MCL subclasses formed a coherent subtree with a single root and strong bootstrap support. However, on the basis of this tree, subclass 2.1 can be clearly subdivided into two subclasses as it included two well supported subtrees (2.1.1 and 2.1.2 in Figure 3). There are several exceptions: subclass 2.3 is embedded in the larger subclass 2.1, suggesting that the 2.1 outgroup is misclassified (grey arc d); and the well-supported subclass 2.2 is framed by divergent sequences originating from the center of the tree (black arcs a and b). In addition, in the same region, a small number of sequences among the deepest branches have an MCL classification in disagreement with their topological location on the tree (black arc c).

**Figure 3:**
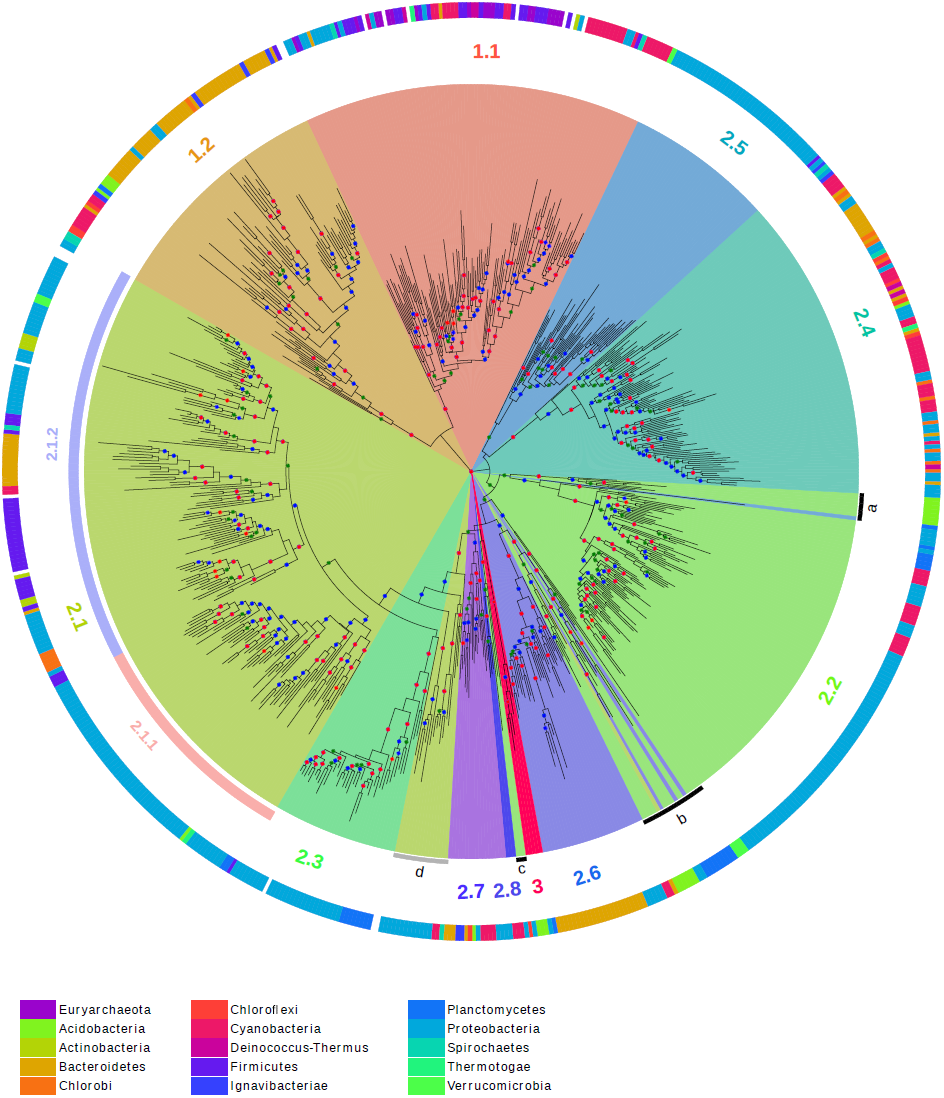
Tree of the TnpA_Y1_ representative sequences. The tree is rooted to the branch that separates class 1 and 2. Color code for the bootstrap value intervals: green in [0.7, 0.8], blue in [0.8, 0.9] and red in [0.9, 1]. The MCL subclasses obtained with IF=3.0 are colored and subclass numbers are added in the corresponding segments. Based on the tree topology, subclass 2.1 can be further divided in groups 2.1.1 and 2.1.2 and three subsets of sequences appear misclassified by MCL (highlighted by black and grey segments) according to their tree location. The taxonomic origin of the sequences is shown as a colored code in the outer ring (NCBI taxonomy phyla level). The phyla with less than three sequences on the tree are not colored. The Phyla are color-coded at the bottom of the figure. The tree display was obtained with online iTOL [83] (https://itol.embl.de/).

### Taxonomic distribution of the subclasses

To determine the taxonomic distribution of the different subfamilies, the sequences of the tree were annotated according to the phylum to which their species belongs by following NCBI taxonomy (Figure 3, outer ring). The subclasses display very different phylum diversity. Subclass 1.1 (TnpA_IS*200*_) exhibits high taxonomic diversity. However the phyla are fragmented into several patches. This patchwork distribution is expected for mobile elements such as IS*200*/IS*605* which undergo horizontal transfer between distantly related phyla. Subclasses 2.1.2, 2.4 and 2.7 also exhibit a similar trend. In contrast, subclasses 2.1.1, 2.3, 2.5, 2.6 and 2.8 are dominated by a single phylum implying a phylum-specific *tnpA*_Y1_ gene expansion. Indeed, sequences from the subclasses 2.1.1, 2.3 and 2.5 are mostly found in Proteobacteria, sequences from the subclass 2.6 in Bacteroidetes and sequences from the subclass 2.8 in Cyanobacteria.

The tree was computed on a subset of our initial sample composed of potentially functional proteins with a complete catalytic core domain. To obtain a global view of subclass distribution across bacterial phyla, we compiled the results from the original sample, retaining only one strain per species. Among the 1,354 species, 57.2% do not encode a *tnpA*_Y1_ gene, 25.6% encode at least one member of class 1 (with 21.5% at least one member of subclass 1.1), 25.4 % encode at least one member of class 2 and 0.3% at least a member of class 3. Multiple different class 2 genes can co-occur in several species. Of these, we have distinguished up to four genes from different class 2 subclasses in 12 of the species in the library. Figure 4 shows the presence or absence of subclass members in each strain. Subclass 1.1 members are well distributed throughout, with a few strongly represented phyla (*e.g.* Cyanobacteria, Clostridia…). Members of the other subclasses show sparse distribution but with a tendency to co-occur in the same genera genome set (Cyanobacteria, Bacteroidetes, Epsilon-, Delta-, Beta- and Gamma-proteobacteria). Subclass 2.8 appears restricted to Cyanobacteria. The assembled Tenericute genomes, represented by 35 Mollicute species in our sample, do not exhibit *tnpA*_Y1_ homologs (although a number of IS*200*/IS*605* can be found in unassembled genomes). Class 3, enlarged to 24 proteins, is principally present in Planctomycetes (11), Acidobacteria (6) and Proteobacteria (5).

**Figure 4:**
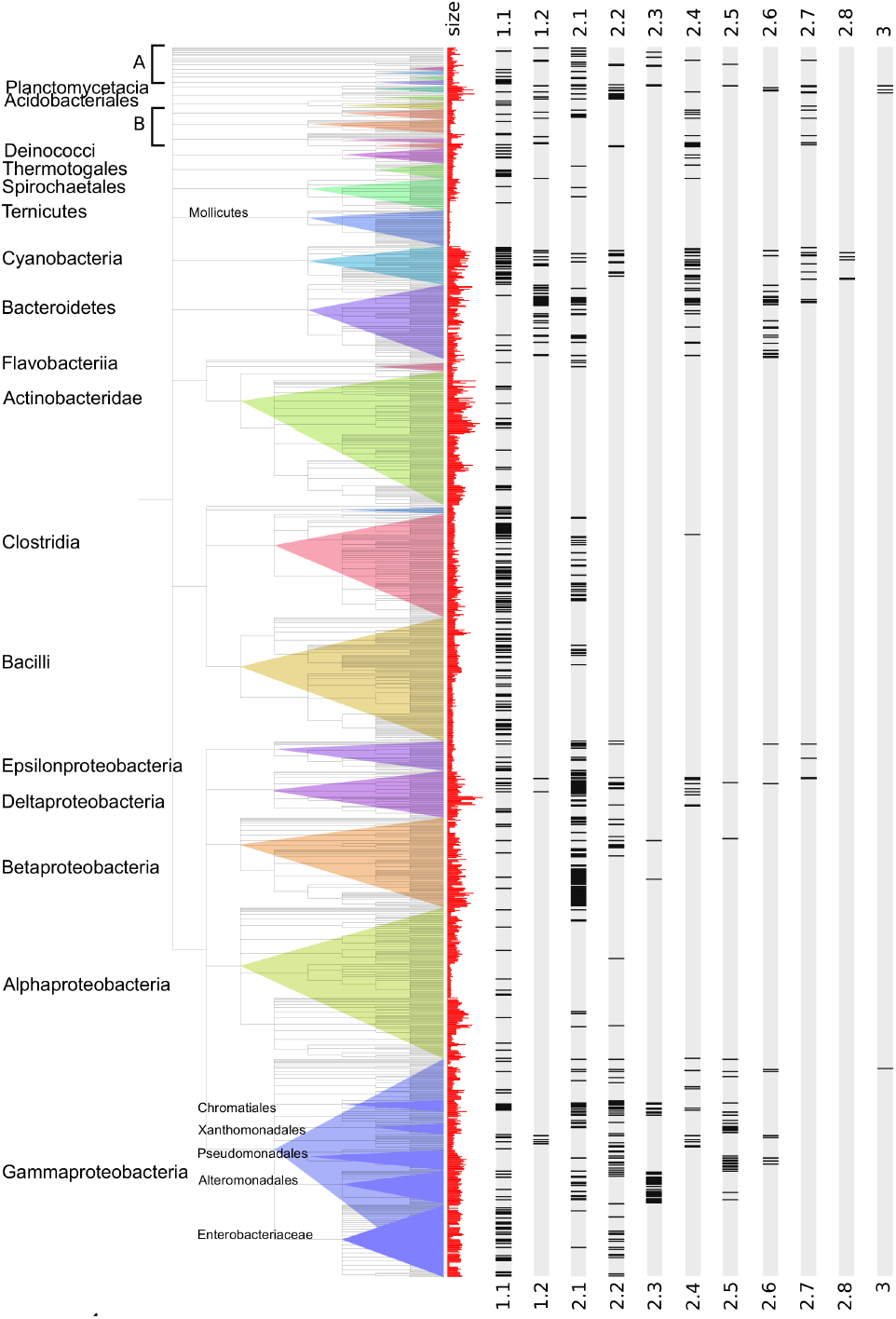
Taxonomic distribution of the TnpA_Y1_ subclasses. Each species of our sample of complete bacterial genomes is represented by one strain and displayed according to the NCBI taxonomic tree (phyla levels). Names of the largest phyla are indicated, first column, histogram of the genome sizes (red), next columns gene occurrence for each subclass. Black bands indicate the presence of at least one copy of the subclass in the genome. A: includes Deferribacterales, Nitrospirales, Synergistia and Fusobacteriales. B: includes Aquificales, Chlorobia, Chlamydiales and Chloroflexi.

In bacteria, genome size variation is associated with the gain and loss of accessory genes [39,40]. To test whether *tnpA*_Y1_ gene distribution follows a similar trend, we compared genomes coding for at least one of these genes with that containing none. We observed that the mean genome size is significantly larger for the former than for the latter (p value < 10^−16^; Figure S5A). This effect is stronger when the TnpA_IS*200*_ of subclass 1.1 are removed (Figure S5B and S5C). Strains with genome sizes greater than 5 Mb have a very high probability of encoding at least one *tnpA*_Y1_ gene. Notable exceptions in our sample are the high G+C Gram-positive Actinobacteridae, that have among the largest genomes but carry only members of subclass 1.1, and Alpha-proteobacteria that include both species with small (such as the Rickettsiales) and large genomes (such as Rhizobiales) but exhibit none or only a few *tnpA*_Y1_ homologs per genome regardless of the genome size. This general preponderance of *tnpA*_Y1_ (excluding the IS-associate proteins) genes in larger genomes suggests that these genes might contribute to the adaptation of their host to new selective pressures and/or to diverse ecological conditions.

### Subclass domain organization

To better define the organization of protein domains, a multiple-alignment of the core domain was performed (Figure 5) using the sequence order from the maximum likelihood tree (Figure 3) and trimmed for sites with rare insertions. This figure shows that core domain length and the distance between the HuH and Y motifs (Figure 5 top) is quite different between families but is conserved within each family. To facilitate further analyses the core domain was split into upstream and downstream sub-domains (annotated in blue; Figure 5 top). The upstream sub-domain ends a few residues after the HuH motif and the downstream sub-domain starts several residues before the conserved W residue. The major length variations are concentrated between the strongly conserved structural elements, β3 and αC and between αC and β4. The insertion domains between β3 and αC are restricted to subclasses 2.3 and 2.4 but differ from each other. The sequence of the 2.3 insertion domain (55 AA) is strongly conserved while the 2.4 insertion domain is highly variable both in length and sequence. Inspection of the DNA sequences of members of this subclass with MEME revealed short repeats that could explain the observed heterogeneity in subclass 2.4 domain (Figure S9).

**Figure 5:**
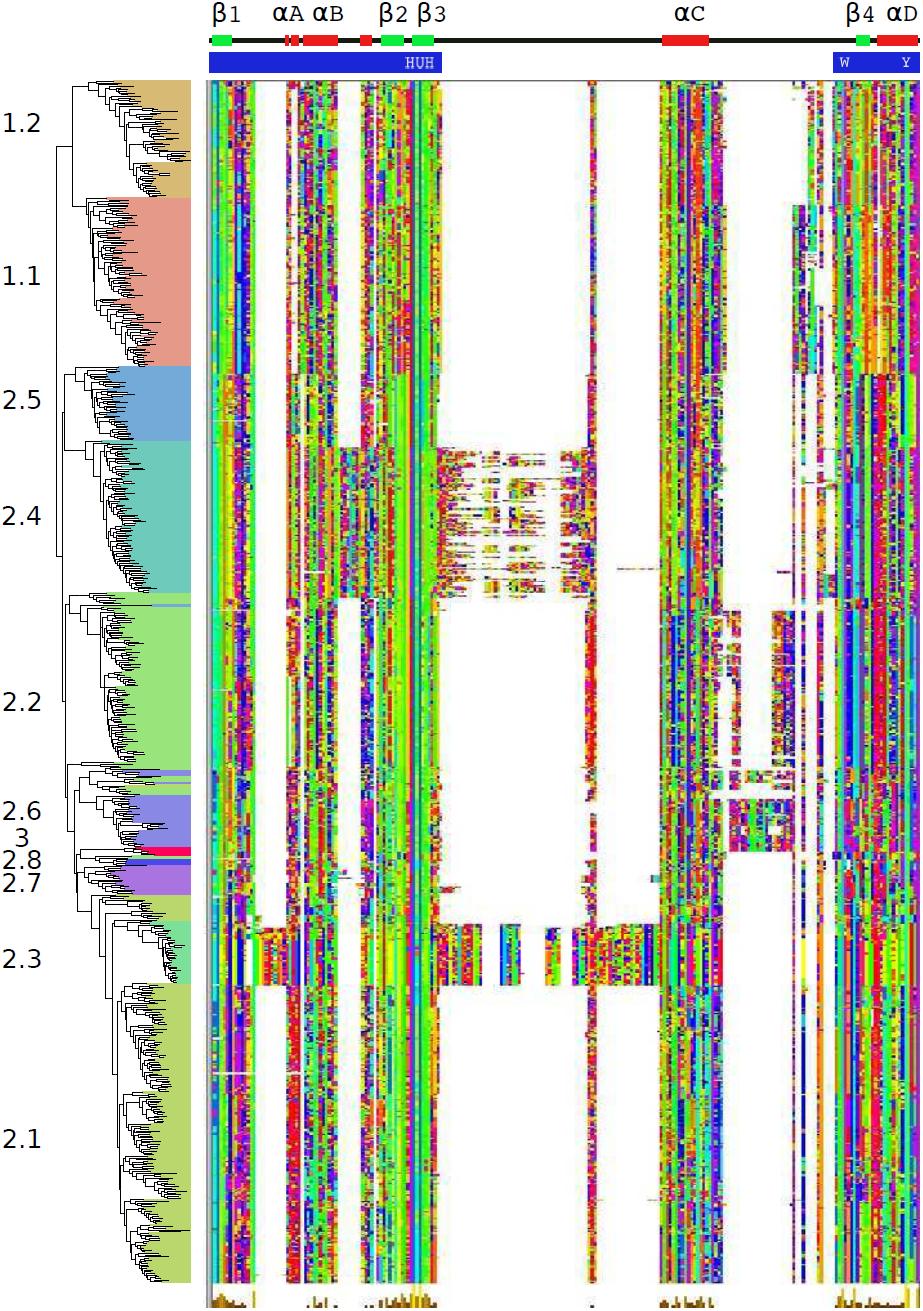
Multiple alignment of the conserved core region. The alignment was reordered according to the phylogenetic tree. The secondary structure of *E. coli* TnpA_REP_ (PDB 4ER8) is shown above the alignment. The location of the core upstream and downstream sub-domains is shown by blue bands. The alignment was trimmed to remove rare insertions and the display was obtained with Jalview and the Taylor color option.

Additional small insertion domains are observed, between β1 and αA (10 AA) in subclass 2.3 and at the end of αB (8 AA) in subclass 2.4. These are located in similar parts of the 3D structure (Figure S6A and S6B respectively). These domains occur far from the projected DNA/protein interaction surface suggesting that they may not be involved in DNA binding or processing. The short insertions observed in subclasses 2.2 (10 AA) and 2.6 (21 AA) between αC and β4 are located close to a structural region involved in REP recognition (Figure S6C-E) suggesting they might contribute to DNA/protein interactions.

To characterize the overall class-specific domain organization and sequence conservation, we designed new HMMER profiles for the full length proteins. The protein domain map was constructed with reference to the phylogenetic tree (Figure 6A). The different proteins are schematized as horizontal grey lines with color segments highlighting the different domains (Figure 6B-D). This reveals that class 1 members are shorter and more homogeneous in size than are class 2 proteins. Note that sequences of subclass 2.1.2 have a higher heterogeneous length distribution.

**Figure 6:**
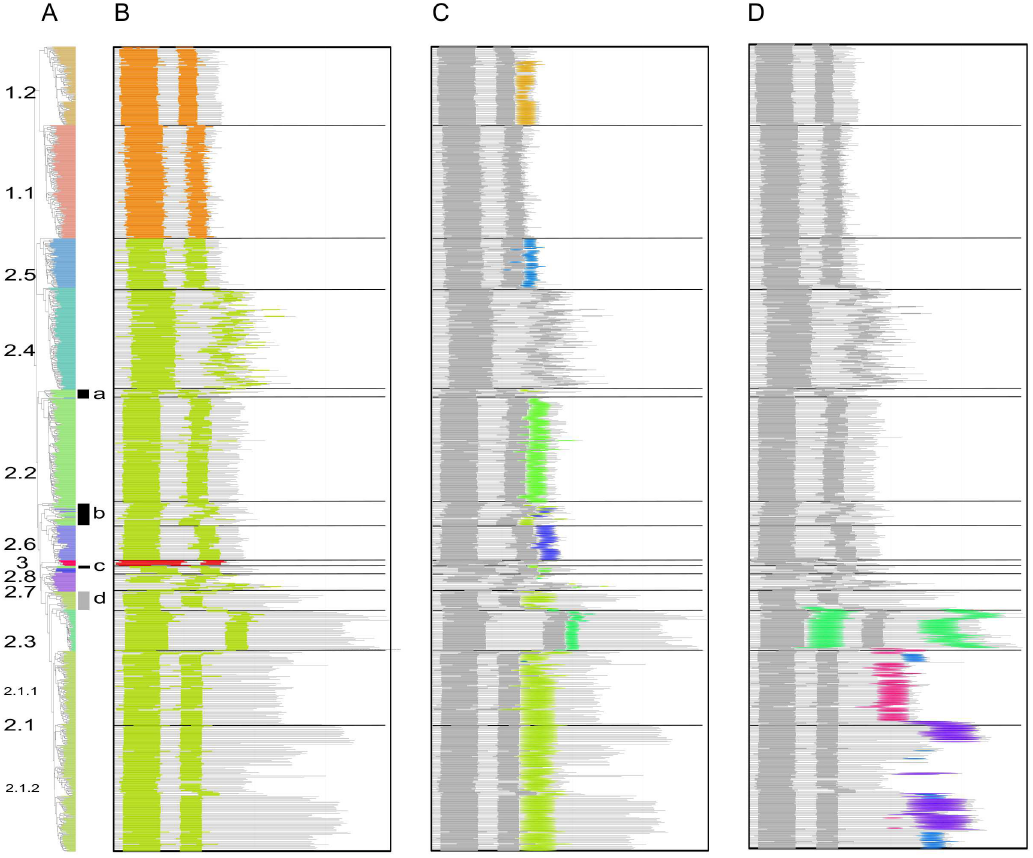
Global domain organization. Proteins are shown as horizontal grey lines with colored patterns that refer to distinct features. Each panel highlights sequential protein domains. A. Phylogenetic tree with subclass annotations. The black and grey segments highlight subsets of sequences misclassified by MCL (Figure 3). B. Positions of the upstream and downstream sub-domains of the core domain are shown in orange for TnpA_IS*200*/IS605,_ green for TnpA_REP_ and red for class 3. C. Short conserved domains found next to the core domain. D. Additional conserved domains found in C-ter regions and between two half of the core domain. E. Intrinsically disordered regions.

In general the distance between conserved core upstream and downstream sub-domains is well preserved within each family although in subclass 2.4 it is quite variable as previously mentioned (Figure 6B). Short length variations between subclasses are observed at the protein N-terminus. These are longer and more variable in the small subclass 2.7. Figure 6C highlights the domain following the core domain. Most classes, except subclass 1.1 and subclasses 2.4 and 2.7, exhibit such a core domain extension with different subclass-specific conserved domains as judged by their HMMER profiles (Figure S7). Moreover, domains of subclasses 2.1, 2.2, 2.3 and 2.6 exhibit similarities in their extension regions (Figure S7) including a conserved Serine. These short motifs were also detected by MEME (Figure S3). The sequences highlighted by the arc a (Figures 3 and 6C) that appear misclassified in subclass 2.2 do not actually share the specific subclass 2.2 extension. The sequences featured by arc b (Figures 3 and 6C) show a non-homogeneous extension with some sequences sharing the subclass 2.6 C-terminus. Finally, the 2.1 proteins placed as an outgroup of subclasses 2.1 and 2.3 (grey arc d, Figures 3 and 6C) possess the subclass 2.1 extension motif.

Additional conserved domains were identified in the longest sequences (Figure 6D). Subclass 2.3 carries a specific C-ter domain (Figure S8A) in addition to the conserved domain (Figure 5) between upstream and downstream core segments (Figure S8A). Division of subclass 2.1 into subgroups 2.1.1 and 2.1.2 suggested by the tree topology (Figure 3) is reinforced by the C-ter organization. Subgroup 2.1.1 carries a conserved C-ter extension, while subgroup 2.1.2 is more heterogeneous both in length and by the presence or absence of conserved sequences (purple and blue, Figure 6D). The 2.1.2 subgroup additional C-ter domains include a region with similarity to Pfam Bac_DnaA_C domain (purple), a motif found in DnaA proteins which binds 9-bp repeats upstream of the oriC replication origin and activates bacterial DNA replication initiation. Other C-ter ends have similarities with either the Pfam HTH_23 or HTH_28 and are likely to be involved in DNA binding. The Bac_DnaA_C (purple) and HTH (blue) related domains (Figure S8B) share conserved residues and may correspond to a long and short version of an ancestral helix-turn-helix domain.

### REP-like sequences in the vicinity of *tnpA*_REP_

Full length *tnpA*_REP_ genes are often flanked by REP sequences present in multiple copies dispersed in intergenic regions of the host genome [5,22,36,37]. Each *tnpA*_REP_ is associated with a specific REP sequence family. A genome may carry more than one type of *tnpA*_REP_ gene but these are all associated with their own specific REP sequence family [22,34,35,37]. A given REP sequence family can include different subfamilies, as observed in *E. coli* MG1655 where three different related REP sequences occur (Y, Z1 and Z2) [31], and its single REPtron is composed of Y and Z2 REPs arranged in three REP sequence pairs in inverted orientation (three REPINs). It has been proposed that TnpA_REP_ is involved in the mobility and/or amplification of REP sequences [5,22,32,33] and the specific nuclease activity of *E. coli* TnpA_REP_ [5,23] strongly supports this hypothesis.

To determine whether the other *tnpA*_REP_-like genes are also associated with REPs, we searched for putative REP sequences in each genome with RNAMotif [41] (Materials and Methods). Canonical REP sequences, with a 5’ GT[AG]G tetranucleotide sequence at the foot of a secondary structure, were found in the close neighborhood (+/−1000 nt) of the large majority of the subclasses 2.2, 2.4, 2.5, 2.7 and 2.8 genes. The detailed results are shown for sequences used to compute the phylogenetic tree (Figure 7C). This program recovered the vast majority of REP sequences which had been identified manually (Figure 7B, red rectangles). However, in a few instances the program did not detect manually identified putative REP sequences (Figure 7B, violet rectangles). For classes 2.6 and 3 a manual inspection was necessary to identify putative motifs since these diverge significantly: they do not carry the consensus GT[AG]G sequence and have a weaker secondary structure (Table S1). In nearly half the sample, as in *E. coli*, each gene was flanked by more than one REP sequence subfamily (Figure 7D). Neither automatic nor manual searches identified REP-like sequences associated with the 2.1 and 2.3 TnpA_Y1_ subclasses.

**Figure 7:**
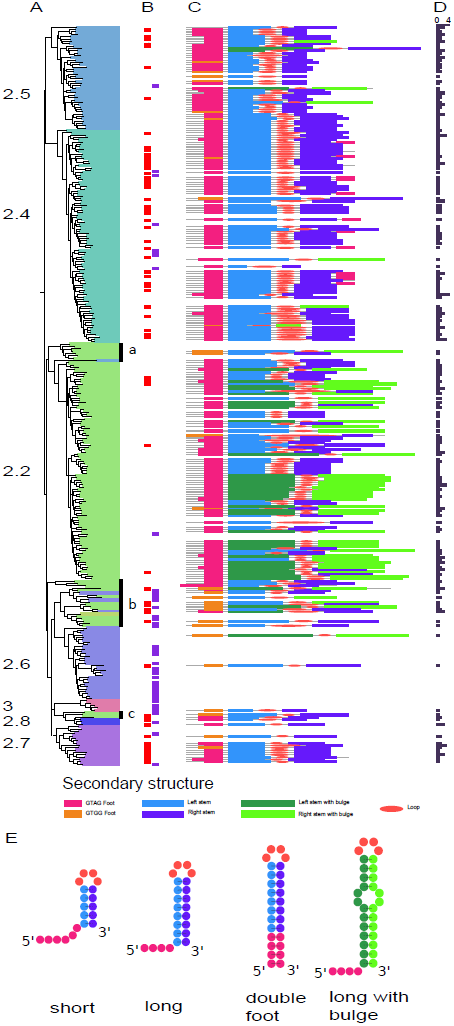
REP secondary structures and distribution. A. Phylogenetic tree with subclass annotations. B, REP sequences annotated by manual inspection: red rectangles, REP sequences also found by automatic annotation and violet rectangles, REP sequences missed by automatic annotation. C, schematic representation of REP sequences found by automatic annotation, only the more frequent REP sequence family is reported for each REPtron. D, Histogram of the number of REP sequences families per REPtron. E, Distinguishable REP sequence secondary structure types.

#### Class specific REP sequence features

REP sequences can have variable lengths with either long or short hairpins (Materials and Methods). The REP sequence stem length distribution is TnpA_Y1_ subclass specific. Subclass 2.2, which contains the *E. coli* TnpA_REP_, includes a majority of REP sequence families with longer stems showing miss-pairing (bulges on the secondary structures, Figure 7E). The *tnpA*_REP_ genes assumed to be misclassified in subclass 2.2 (Figure 5 and Figure 6, black arcs a, b and c) are enriched in a GTGG motif confirming their marginality (Figure 7A black bars, a, b and c).

Subclass 2.4 contains two REP types: canonical long REP sequences, where the bottom of the hairpin stem is composed of 3 consecutive G-C base pairs, and particular long REP sequence families with a 5’ GT[AG]G and its reverse complement, C[TC]AC 3’, at the end of the sequence (“double foot”, Figure 7E). Note, that a double foot REP sequence could be generated simply by central deletion between an inverted REP sequence dimer. A unique feature of subclass 2.4 is that, in addition to the REP sequences flanking the *tnpA*_REP_ genes (Figure S10), REP sequences can be found inserted within the *tnpA*_REP_ genes themselves. Remarkably, these insertions do not disrupt the reading frame (Figures 5 and 6). At the protein level, the insertions are generally located between the HuH and Y motifs. REP sequence insertions in a variety of unrelated genes have also been reported. These insertions may not affect the activity of the corresponding proteins if they are located in flexible linkers or loops [36]. On the predictive structural model of subclass 2.4 (Figure S6B), the variable region appears as a peripheral extension away from the ssDNA binding and catalytic sites which may have little impact on protein activity.

Subclass 2.5 TnpA_Y1_ are associated with short REP sequences. In some of these, the 5’ conserved tetranucleotide is located 1-2 nt from the secondary structure rather than directly at its foot but are “compensated” by shorter stems. Thus, the sum of the foot and stem length is distributed in a narrow range ([10-13] nt). In *Stenotrophomonas maltophilia*, a subtree of subclass 2.5, REP sequences are more similar to the long canonical REP sequences of subclass 2.4. TnpA_Y1_ of subclass 2.6 are flanked by atypical secondary structures identifiable by manual inspection (Figure 7). Copies of these secondary structures can be found within the host genome. Members of subclasses 2.7 and 2.8 are flanked by long canonical REP sequences with 5’ GT[AG]G tetranucleotides.

## Subclass specific features summary

### Class 1

Class 1 composed of subclasses, 1.1 and 1.2, is present in 25.6% of our sample of species, close to the 26% estimated previously [38].

#### Subclass 1.1

Subclass 1.1 is the largest group, present in 22.4% of all species (Table 1) and includes typical IS*200* tranposases (ISfinder [11,42,43]). It shows the highest taxonomic diversity of all TnpA_Y1_ proteins (Figure 3 and 4) and it is the only subclass observed in Archaea. A characteristic signature is the Y motif (Y---Q) (Figure S3) and the hydrophobic residue (u) of the HuH motif is either V or I. These are the shortest TnpA_Y1_ family members, composed essentially of the core without additional domains. These *tnpA* genes are often found in microsynteny with a *tnpB* gene whose function is not yet entirely clear [16] and flanked by the typical small subterminal DNA hairpin structures (ISfinder). The high genomic copy number (Figure S11), the presence of partial sequences, the high sequence conservation observed between copies within the same genome (Figure S12) and their wide distribution in Archaeal and Bacterial phyla are consistent with the IS nature of these sequences [11,44,45].

**Table 1:**
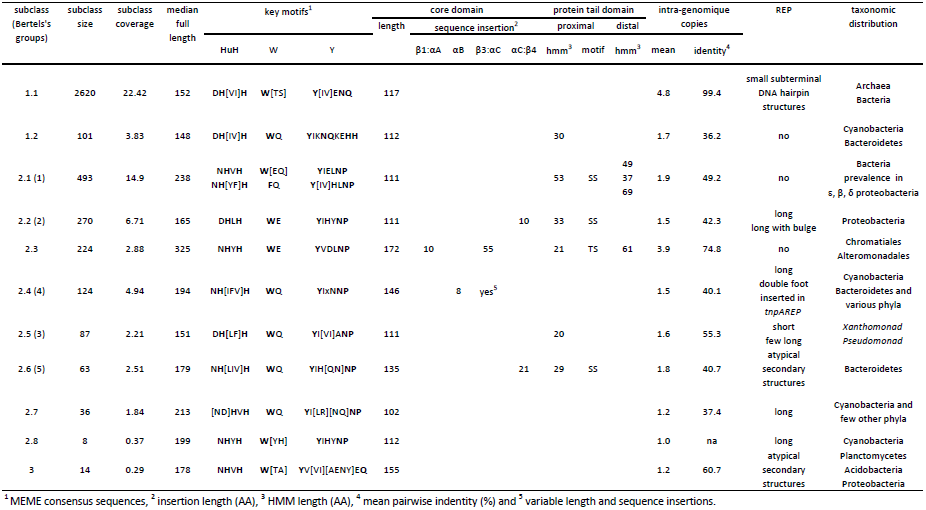
Summary of the TnpA_Y1_ family subclasses properties

#### Subclass 1.2

Subclass 1.2 proteins are closely related to those of subclass 1.1 (Table 1; Figures 2 and 3) but with distinctive features: a characteristic Y motif signature, Y---Q--HH (Figure S3); a shorter core domain (Figure S4), a conserved specific C-ter domain (Figure 6C); a low genomic copy number (Figure S11); generally without typical terminal IS*200* secondary structure features; and a taxonomic distribution mostly restricted to species of Cyanobacteria and Bacteroidetes phyla (covering less than 4% of all species). Therefore, despite their close relationship, this subclass does not share the IS properties of subclass 1.1. It was not described by Bertels *et al.* [38] and to our knowledge, this is its first characterization.

### Class 2

Class 2, present in 25.5% of our species sample, includes the known TnpA_REP_ proteins and is composed of 8 subclasses 2.1-2.8. Class 2 genes have low intra-genomic copy number (except subclass 2.3, Figure S11) and low sequence conservation between copies within the same genome (Figure S12). The common Y-motif signature of all class 2 members is Y---NP. The hydrophobic residue (u) of the HuH motif is either L, V, I, F, Y or M (ranked by decreasing frequencies) (Figure S3).

#### Subclass 2.1

Subclass 2.1 is the second largest subclass (14.9% of all strains, Table 1). Their members are among the longest proteins of our sample. They carry additional C-ter conserved domain (Figure 6 and S8B). However this subclass can be subdivided into two groups, 2.1.1 and 2.1.2, according both to the tree topology (Figure 3) and the C-ter domain nature that appears conserved in the 2.1.1 members while it is more heterogeneous in 2.1.2 sequences (Figure 6D). This C-ter extension shows similarities with helix-turn-helix motifs (Bac_DnaA_C, HTH_23 or HTH_28 Pfam profiles, Figure S8B). They share few conserved residues and may correspond to a long and short version of an ancestral helix-turn-helix domain. Neither automatic nor manual searches identified REP-like sequences in the vicinity of the *tnpA*_REP_ gene. Previously, a collection of Y1 proteins homologous to TnpA_REP_, was called TIRYT (Terminal Inverted Repeat associated tYrosine Transposase) because they were associated with terminal inverted repeats rather than typical secondary structures [36]. Of the 26 examples described [36], 20 belong to subclass 2.1. All include one of the three HTH domains. However, the consensus sequence GGGG[AT][CG]A[CG] observed in the majority of the flanking inverted repeats does not resemble the consensus TT[AT]TNCACA of the high affinity DnaA boxes [46] suggesting that these C-ter domains may not be directly involved in DNA target recognition or have evolved to recognize a new target sequence. The other 6 TIRYT examples are distributed between subclasses 2.2 (3 from arc b and one from arc a) and 2.5 (2 proteins). The TIRs flanking subclass 2.5 *tnpA*_REP_ in *P. putida* and *P. fluorescens* originated from fusion of degenerate REP elements with unrelated sequences. Subclass 2.1 therefore probably corresponds to the TIRYT described by De Nocera *et al.* [36]. Further analyses are necessary to confirm this.

#### Subclass 2.2

Subclass 2.2 includes the first *tnpA*_REP_ identified in *E. coli* [5,22,36] and represents the third largest of the groups (6.7% of all strains). They are predominantly found in Proteobacteria (Figures 3 and 4; Table 1). Members exhibit short insertions between αC and β4, a region neighboring that involved in REP sequence hairpin recognition in the crystal structure [23] (Figure S6D and S6E). It is noteworthy that this subclass includes REP sequences with a hairpin-bulge-hairpin secondary structure (Figure 7) suggesting that these short conserved domains might contribute to an original REP/protein interaction.

#### Subclass 2.3

The members of subclass 2.3 are found in less than 3% of all strains but prevail in Chromatiales and Alteromonadales where more than five copies per genome can be identified (Figure S11; Table 1). However, those copies do not exhibit the high intra genomic sequence conservation observed for members of subclass 1.1 (Figure S12). The subtree corresponding to this class is deeply rooted in subclass 2.1 (long branches) (Figure 3). However, the leaves corresponding to the Proteobacteria are linked by short branches which suggest a recent and active spreading of elements of this distinct TnpA_Y1_ subclass in Chromatiales and Alteromonadales strains. As for subclass 2.1, the gene is not flanked by identifiable REP-like DNA secondary structures. Members have the longest sequence length (median equal to 325 AA; Figure S1) resulting from multiple domain additions. Two domain insertions occurred between conserved secondary structure elements of the conserved core region (β1 and αA; β3 and αC). These domains are highly conserved and exposed to the surface of the protein (Figure S6A). Moreover, the long C-terminal tail is composed of two domains. The proximal domain includes a conserved serine residue observed in other subclasses while the long distal domain is subclass specific. The accessibility of all these additional domains suggests that they may be involved in molecular interactions and/or processes not yet identified.

#### Subclass 2.4

Subclass 2.4 is abundant in Cyanobacteria and Bacteroidetes but also scattered across a number of phyla (Figures 3 and 4; Table 1). Subclass 2.4 contains two REP types: canonical long and double foot REP sequences. A remarkable and unique feature of subclass 2.4 members is the predominance of REP sequence insertions within the *tnpA*_REP_ genes.

#### Subclass 2.5

The cognate REPs of subclass 2.5 belong to the larger REP sequence families with typical REP sequence organization. Elements of this subclass are mostly limited to *Xanthomonad* and *Pseudomonad* species (Figures 3 and 4; Table 1) and some have been described in previous studies [32–34,47,48,48–50]. Comparative genome analysis suggests a recent invasion of *tnpA*_REP_ genes in these genomes and that REP sequence diversification and proliferation are still ongoing leading to high numbers of REP copies [37]. All these observations are in agreement with the suggestion that *tnpA*_REP_ genes coevolve with their cognate REPs [22].

#### Subclass 2.6

Subclass 2.6 members are mostly present in Bacteroidetes (Figures 3 and 4; Table 1). They are flanked by atypical secondary structures identifiable by manual inspection (Figure 7), copies of which can be found elsewhere in the genome. Like subclass 2.2, they also contain short insertions between αC and β4 (Figure S6D and S6E). This domain may contribute to the recognition of the unusual REP sequences.

#### Subclasses 2.7 and 2.8

Members of small subclasses 2.7 and 2.8 are variable in length (Figure 6; Table 1) and genes are flanked by long canonical REP sequences with 5’ GT[AG]G tetranucleotides. Subclass 2.8 is restricted to Cyanobacteria in our sample but closely related sequences are found from other unrelated phyla in NCBI non-redundant (nr) database. They are characterized by a conserved Y residue in the HuH motif (HYH) and a proline rich N-ter domain (Figure S3). Some carry a REP sequence insertion.

### Class 3

Class 3 proteins are relatively rare (<0.3% of all species) and found principally in the Planctomycetes, Acidobacterial and Proteobacterial phyla (Table 1). They include a Y-motif signature, Y Q related to that of class 1 (Figure S3), and a distinctive core domain profile (Figure 6B). They are flanked by several atypical secondary structures with a 5’ tetranucleotide foot CnGA identifiable by manual inspection (Figure 7, Table S1). This feature is more reminiscent of a REPtron than an IS. Moreover, the associated DNA secondary structures can be present in high copy number in a given host genome (e.g. ~ 600 in *Solibacter usitatus* Ellin6076), again reminiscent of REP sequences. Interestingly, there are three closely related class 3 genes in this bacterium. Thus class 3 examples resemble both IS and REPtrons and are possibly evolutionary intermediates between class 1 and 2 (Figure 2C). Further experimental analyses are required to determine their origin and behaviour.

### Subclass partition: strength and weakness of different approaches

Bertels *et al.* (2017) defined six TnpA_Y1_ groups, with at least 30 or more members, for both TnpA_IS*200*_ and TnpA_REP_ (RAYTs) families. The structure of these two families appeared very different. The TnpA_IS*200*_ family was more homogeneous and the groups (clusters) were very close to each other while those of the TnpA_REP_ (RAYTs) family were more distinct. Our analysis structures the TnpA_REP_ (RAYTs) family into eight groups. To establish the correspondence between both classifications we used the sequences provided by Bertels and collaborators (supplementary data) for their five larger RAYT groups. The IS group sequences and the RAYT group 6 sequences were not available. 291 of the 292 RAYT group 1 sequences are assigned to our subclass 2.1; 238 of the 239 RAYT group 2 sequences belong to our subclass 2.2; 113 of the 120 RAYT group 3 sequences are allocated to our subclass 2.5; the 100 RAYT group 4 sequences are assigned to our subclass 2.4 and finally the 40 RAYT group 5 sequences are found in our subclass 2.6. Our classification is therefore only partially covered by the groups defined in Bertels *et al.* (2017). One of our larger subclasses (2.3) and the smallest (2.7, 2.8) were not identified. Neither did they detect class 3, whose characteristics suggest that it may represent an evolutionary intermediary between classes 1 and 2. In addition, the well supported distinction between TnpA_IS*200*_ subclass 1.1 and the related subclass 1.2 was not found and the evolutionary relationships between TnpA_IS*200*_ and TnpA_REP_ (RAYTs) families was not clearly established.

These differences could be explained by the different methodological workflows implemented. First for the identification of the TnpA_REP_ (RAYT) and TnpA_IS*200*_ protein candidates, Bertels *et al.* (2017) used two independent tblastn searches based on two queries, one for each protein family (TnpA_IS*200*_ and TnpA_REP_) while we undertook a more sensitive profile based search (hmmersearch) using the transposase IS*200* like (Y1_Tnp) profile from Pfam. At this step, an e-value threshold of 0.01 was used to maximize detection of remote homologs. Sequences of subclasses 2.3, 2.7, 2.8 and 3 could be too distant from the Bertels query sequence (*Pseudomonas fluorescens* SBW25, PFLU_RS20900 belonging to subclass 2.5) to be identified by their tblastn approach (e-value < 0.02). In addition, we then reduced the sequence redundancy due to the recent gene duplications and the unbalanced representation of bacterial strains in the dataset by clustering closely related sequences (cut off of 70% of identity), each cluster being further reduced to one representative sequence, the medoid.

Exploration of sub-family organization was performed in both analyses using MCL graph partitioning but the similarity relationships between pairwise sequences were calculated differently. We generated a reciprocal all-against-all BlastP comparison (e-value < 1e^−5^) on our data set including both TnpA_IS*200*_ and TnpA_REP_ (RAYTs) proteins and the BlastP results were reformatted to an undirected and weighted graph (log of e-value). Different MCL inflate factor values were tested to select the optimal partition. Our approach is inspired by TRIBE-MCL [51] (https://micans.org/mcl/), a method for detecting protein families in large databases. Bertels *et al.* (2017) computed relationships between proteins of the TnpA_IS*200*_ or TnpA_REP_ (RAYTs) family independently by pairwise sequence comparisons obtained with a global alignment algorithm (Needleman–Wunsch). The results were filtered to retain only the pairs with more than 26% identity and then converted into a weighted graph (pairwise identity). The use of such a global alignment does not appear appropriate since, as we describe above, the proteins show variable domain organization and length heterogeneity (Table 1). To explore the relationships between the groups identified independently in both families they then randomly selected 30 sequences in each group and partitioned the graph using MCL as explained above. An inflate factor of 2.0 was invariantly used for graph partitioning with MCL.

Finally, we explored the evolutionary relationships between the proteins of the family by constructing a tree based on the conserved core domains shared by the entire sequence set. Only potentially functional sequences (with the conserved HuH and Y motifs) were retained because pseudogenes with a higher evolution rate could disturb tree topology. Bertels *et al.* (2017) computed a phylogenetic tree by randomly selecting three sequences from each of the TnpA_IS*200*_ and RAYT groups.

### Vertical versus horizontal inheritance

Although there is no evidence that class 2 genes are mobile, their presence in distantly related species suggests that the ancestor appeared early in evolution. However, members of this class are observed in only 25.5% of species of our sample with a patchy taxonomic distribution, in agreement with Bertels *et al.* (2017). The hypothesis of vertical inheritance would imply multiple independent gene loss events during the course of evolution.

A more parsimonious model is to assume that dissemination of class 2 genes occurred by multiple horizontal gene transfers (HGTs). Support for this hypothesis comes from the observation of *tnpA*_REP_ gene transfer from a Pseudomonad to marine gammaproteobacteria [22] and evidence that HGT is likely to have occurred between fluorescent pseudomonad strains [37]. In the absence of formal evidence for autonomous mobility of these elements, *tnpA*_REP_ genes might use the same routes as accessory genes for transfer (e.g. transformation [52], conjugation [53], transduction [54], and gene transfer agents [55,56]). The taxonomic distribution of class 2 genes (Figure 4) is consistent with the observation that HGT events are more frequent between closely related species [57]. Indeed, except for subclass 2.1 and to a lesser extent for subclass 2.4, *tnpA*_REP_ genes are mostly confined to a specific taxonomic group. This proximity would also favour integration of class 2 proteins into host cell metabolism by, for example, providing binding sites for host proteins or influencing gene expression. The observation here, that intra-genome gene copies belonging to the same subclass are distantly related (Figure S12), is also compatible with multiple HGTs. Gene loss should also be considered as an evolutionary force since such events are suspected in genomes encoding REP sequences in the absence of *tnpA*_REP_ gene [36] and is well documented in *E. coli* strains, where the REPtron locus had been replaced by an operon encoding toxin/antitoxin genes [23]. Gene loss may have occurred after the acquisition of *tnpA*_REP_ by HGT in the ancestor. The contribution of HGT and gene loss can be estimated with reconciliation methods that compare gene trees and species trees to recover the history of gene families [58], but this is outside the scope of this work.

### Functional outcomes

All sequences share a conserved structural core domain including HuH and Y motifs involved in the distinctive site-specific ssDNA cleavage and ligation mechanism of this protein superfamily [10,15,23]. However, the local context of the catalytic tyrosine is class-specific: Y---Q, Y---NP and Y Q for classes 1, 2 and 3, respectively. The position of the conserved glutamine residue (Q131 in the *Helicobacter pylori* IS*608* transposase, Figure 1), on the same face of αD helix as the catalytic tyrosine residue, is essential for transposition *in vivo* and this residue is part of the divalent metal ion binding site with the histidines of the HuH motif [10]. The same role was assigned to the asparagine residue (N119) in TnpA_REP_ from *E. coli* MG1655 [23]. The sequence conservation observed suggests that proteins of different classes have the same single-stranded DNA editing activities.

The Y1 transposases including TnpA_IS*200*_ and TnpA_REP_ of *E. coli* require ssDNA. Although this can be provided by a variety of different processes [16,17,21,23,59], the lagging strand of the replication fork is an important source *in vivo* [60]. The propensity for excision from and insertion into the lagging strand and the intimate coupling of single strand transposition to replication has been well documented for two members of the IS*200*/IS*605* family [11–13,16,19].

Transposase targeting can be mediated by direct interaction with proteins of the replication fork complex. For example, TnpA of IS*608* (TnpA_IS*608*_) and other TnpA_IS*200*_ family members are thought to interact with DnaN [21,61], the β sliding clamp replisome processing factor. Moreover, TnpA_IS*608*_ shows affinity for DNA structures resembling replication forks [21]. It has been proposed that the region of TnpA_IS*608*_ interaction with DnaN is located in the C-ter region, next to the catalytic tyrosine, in the αE helix [61]. A survey of enzymes that interact with β sliding clamp within the replication fork reveals that they share a short and poorly conserved binding motif (consensus QL[SD]LF) [62]. These motifs are often located in a highly flexible C-terminal tail of the protein [62–64]. However, this sequence is not conserved within TnpA_IS*200*_ members. In spite of this, it seems possible that the short serine motif located next to the catalytic tyrosine in subclasses 2.1, 2.2, 2.3, and 2.6 could play a role of targeting the protein to the replication fork *via* protein-protein interactions. Other subclasses may have evolved alternative subclass-specific motifs or domains for interaction with other host targets. For example, the DnaA-like DNA binding domain in subclass 2.1 proteins suggests an alternative, more direct, way of targeting the replication fork. Interaction of transposase with universal and highly conserved proteins such as sliding clamps would facilitate transfer between phylogenetically distant organisms [65]. Indeed, the transposition pathways of a number of IS and transposons require host enzymatic functions [65–68], including DNA polymerases and other factors implicated in DNA replication, suggesting a functional link between transposition and replication [17,21].

The discontinuous taxonomic distribution of class 2 proteins supports the notion of a transitory positive impact of REPtrons on host cell fitness. Although their function has yet to be formally elucidated, their possible involvement in single-stranded DNA editing and in the proliferation of small dispersed repeated sequences could be a beneficial factor in adaptive transition phases. After this phase, gene silencing or gene loss could restore genome stability. Their phylogenetic proximity to TnpA_IS*200*_ suggests that class 2 proteins may represent mobile element domestication [5,22,32,38].

## Conclusions

Previous studies have underlined the similarities between IS*200*/IS*605* family transposons, their HuH-Y1 transposases and their terminal DNA secondary structures, and the bacterial REP systems, the TnpA_REP_ (RAYT) proteins and their associated REP sequences [5,22,23]. Here, we performed a genome-wide analysis of TnpA proteins related to those of the IS*200*/IS*605* and REPtron families in complete genomes of archaea and bacteria. We refer to these collectively as TnpA_Y1_. All sequences share a conserved structural core domain including HuH and Y motifs involved in the distinctive site-specific ssDNA cleavage and ligation mechanism of this protein superfamily [10,15,23]. Based on sequence similarity, proteins can be arranged in classes and subclasses. Subclass 1.1 includes sequences similar to IS*200*/IS*605* transposases while proteins of the other subclasses are not. Here, we identify and characterize a new subclass (subclass 1.2) closely related to IS*200*/IS*605* transposases but which does not share IS properties, as well as a previously unknown group of HuH-Y proteins (class 3). Each subclass is characterized by specific additional sequence domains possibly involved in protein/DNA or protein/protein interactions and in targeting these proteins to single-stranded DNA. The taxonomic distribution reveals that more than 25% of the analyzed species encode at least one *tnpA*_REP_-like gene, but with accumulation in some phyla. The average size of genomes containing at least one *tnpA*_REP_-like gene is significantly larger than that of genomes without any of these genes. Their patchy taxonomic distribution is in agreement with dissemination by multiple horizontal gene transfers followed by gene loss. The genes, of a closely related subset of subclasses, are flanked by typical REP sequences in a structure called REPtron. The proteins encoded by these genes are assumed to be responsible for REP sequence proliferation in the genome. Proteins of all subclasses share a common catalytic domain involved in single strand DNA editing. The disparity observed in taxonomic distribution and the diversity of domain arrangements of the proteins belonging to different subclasses suggest that they evolved to different cell physiology and raise the possibility of their domestication in cell function related to single strand DNA edition.

## Materials and Methods

### Building TnpA_Y1_ dataset

Completely sequenced genomes of 172 archaea and 2,178 bacteria covering 1,351 distinct species were downloaded from EBI (http://www.ebi.ac.uk/genomes/). The complete genomes of these 2,355 strains, their proteomes and EMBL features were managed with an in house mySQL database. We also downloaded the collection of hidden Markov models (HMM) from Pfam (http://pfam.xfam.org, release 29.0).

To retrieve transposase IS*200*-like proteins, we adopted a two step approach. First, the Pfam entry Y1_Tnp (PF01797), covering the Transposase IS*200*-like protein family, was used as query with hmmersearch from the HMMER3 package (http://hmmer.org/; [69]) against the 7,223,104 protein sequences from our sample. Only proteins showing an alignment with an e-values < 0.01 (recommended threshold) were retained. A first set of 4,081 proteins was obtained. Second, we reduced the sequence redundancy inherent to our sample due to i) the multiple replicates from strains of the same species (e.g. *E. coli* includes 56 strains in our sample), ii) the unbalanced distribution of bacterial phyla in public databases, leading to the overrepresentation of some closely related species and iii) finally the presence of a high number of identical copies in some genomes due to the repetitive nature of the element (e.g. 99 copies of IS*200* in *Yersinia pestis* Angola). To minimize this bias which can affect the performances of multiple alignment and phylogenetic methods, closely related sequences were represented by one sequence, the medoid. To identify this, we performed all-against-all BlastP comparisons of our initial set of proteins. An identity cut off of 70% was chosen for retaining similarities as it offers the best compromise between the sample size reduction and the conservation of the overall TnpA_Y1_ family structure. Protein relationships were then converted into a graph in which the vertices represent protein sequences, and the edges represent their relationships. The graph was further processed by a graph-partitioning approach based on the Markov Clustering algorithm (MCL, [70]). This identified 687 clusters composed of a unique sequence and 244 clusters composed of closely related sequences. For each of the 244 clusters, we computed the medoid, *i.e.*, the sequence whose average dissimilarity to all the other proteins in the cluster is minimal, for which we add the constraint that its length should be close the median length of all sequences of the cluster. This resulted in a set of 931 proteins composed of 687 unique sequences and 244 medoids. Finally, all-against-all BlastP comparisons (e-value < 1e^−5^) of the restricted 931 protein sequence set were performed. The graph obtained was composed of a large connected component of 924 vertices and of singletons or two vertices. Pfam protein annotation revealed that, contrary to the 924 proteins, those belonging to the small clusters had best hits with conserved domains unrelated to the TnpA_Y1_ Pfam domain. They were considered as false positives and removed from our sample to yield a final sample of 924 proteins for further analyses.

### HMM profiles

To characterize the different protein families and domain organization, we used the HMMER package (http://hmmer.org). This implements methods using probabilistic models, profile hidden Markov models (profile HMMs), that assign a position-specific scoring system for substitutions, insertions, and deletions [71–73]. This is used for construction of protein domain models and is the basis of the protein domain model database, Pfam [74,75]. As seeds, we used the representative sequences described above. Alignments were obtained with mafft version 7 [76] with default parameters “except --localpair --maxiterate 1000” parameters to ensure a high accuracy. The multiple alignments were edited with Jalview [77] and the core domain was retained following deletion of left and right flanking regions. Partial or highly divergent sequences were removed. The HMMER profiles were built with hmmbuild from the input multiple alignments and assembled in an HMM database with hmmpress. Protein sequences were scanned against this database with hmmscan. The domain annotation file was parsed with a Perl program to select the best non-overlapping domains in each sequence. The residues which aligned with the conserved motifs in the HMMER profiles were extracted and the results of the domain annotation saved in the database.

### Phylogenetic tree

Multiple alignments were processed with trimAl [78] to eliminate aligned positions with a high frequency of gaps. ProtTest [79] was used to select the optimal parameter combination. The resulting parameters included the LG model of sequence evolution with γ-correction (four categories of evolutionary rates), shape parameter and proportion of invariant sites estimated from the data. Phylogenetic trees were computed with PhyML [80,81]. Branch supports were estimated with parametric bootstrap analysis. Trees were drawn and annotated with the interactive Tree Of Life web server (iTOL, http://itol.embl.de/index.shtm) [82,83].

### Motif identification

The search for conserved motifs used MEME program (Multiple Em for Motif Elicitation, http://meme.nbcr.net/meme) [84]. Individual MEME motifs do not contain gaps. The unaligned sequences were submitted to MEME with a motif width between 3 and 10 amino acids. We searched for a maximum of 20 motifs with an occurrence of zero or one motif per sequence since we did not expect that each motif would be present in all sequences. Repeats were predicted in *tnpA*_REP_ genes with ‘any number of repetitions’ option of MEME. The results of MEME were used by MAST (Motif Alignment & Search Tool) program to annotate the motifs in sequences of our database.

### REP sequence prediction

REPs are not conserved in sequence between species (species-specific) and within a genome copies change slightly in sequence and length (imperfect/partial repeats). However, they share a common structural feature: a tetranucleotide guide sequence located 5’ to the foot of a short hairpin. We used the RNAMotif program [41] to search for REP sequence candidates in TnpA_REP_-encoding genomes. This program requires a file that describes the secondary structure interactions and the set of rules used to filter matching sequences which was constructed from the analyses of REPs reported in [36]. It includes 66 previously described REP sequence families found mostly in Proteobacteria. All include a GTAG or GTGG tetranucleotide located between 0 to 2 nt 5’ from a secondary structure and include a basal stem of fully paired 7 to 20 nt and a loop of 2 to 5 nt (short hairpins) or an additional fully paired stem of 1 to 10 nt connected to the first by a bulge of 1 to 3 nt (long hairpins) (see Figure 7E). In addition, the first four basal stem nucleotides include at least three G or C bases (short hairpins) or the GC content of the stem is >= 50% (long hairpins). Only four REP sequences out of 66 do not follow these rules. The RNAMotif program was applied. The REP sequence candidates predicted in DNA sequences of complete genomes by RNAMotif were clustered into families with BlastN and MCL [70]. For each family a multiple alignment was obtained with mafft [76] and the secondary structure computed with RNAalifold [85]. Only families with at least one occurrence located in the vicinity (1000 nt upstream and downstream) of a *tnpA*_REP_ gene were retained. The predicted REPs were assembled in dimers (BIMEs) or clusters, if the distance between consecutive REPs on the genome is less or equal to 150 nt. This *in silico* analysis was supervised by human expertise.

## List of abbreviations

AA: Amino Acids
nt: nucleotides
N-ter: start of a protein or polypeptide
C-ter: end of a protein or polypeptide
REP: Repeated Extragenic Palindromes
BIME: Bacterial Interspersed Mosaic Element
RAYT: REP-Associated tYrosine Tansposases
TIRYT: Terminal Inverted Repeat associated tYrosine Transposase
REPIN: REP doublets forming hairpIN
HGT: Horizontal Gene Transfer

## Declarations

### Ethics approval and consent to participate

Not applicable

### Consent for publication

Not applicable

### Availability of data and material

The datasets used and/or analyzed during the current study are available from the corresponding authors on reasonable request.

### Authors’ contributions

Y.Q., P.S., and M.C. made substantial contributions to conception and design, or acquisition of data, or analysis and interpretation of data. Y.Q., P.S., M.C. and G.F. have been involved in drafting the manuscript or revising it critically for important intellectual content.

## Acknowledgements

The authors thank Bao Ton-Hoang for discussions.

## Authors’ information

